# Learning of speech categories in humans and Zebra Finches

**DOI:** 10.1101/077321

**Authors:** D. Botskaris, B. Kriengwatana, C. ten Cate

## Abstract

The survival of organisms depends highly on their ability to adjust their behavior according to proper categorizations of various events. More than one strategy can be used in categorization. One is the Rule-Based (RB) strategy and the other is Information-Integration (II) strategy. In this research we analyzed the differences between avian and human cognition. Twelve Greek listeners and four Zebra finches were tested in speech category learning tasks. In particular, both humans and Zebra finches had to categorize between Dutch vowels that differ on duration, frequency or both depending on the condition. Feedback was given for correct and incorrect responses. The results showed that humans and Zebra finches are probably using the same methods of learning depending on the categorization tasks that they are exposed to. If Zebra Finches are actually able to acquire (RB) and (II) category structures using the same strategies as humans, the utility of multiple systems of categorization might not be restricted to primates as current literature suggest.

## Introduction

Organisms have to discriminate and categorize between various auditory and visual stimuli in order to adapt their behavior in respect to their environmental conditions. Sometimes, categorization is so crucial that can determine the difference between life and death. As an example, imagine a gazelle being unable to distinguish between the sound of a predator and a mate call. Generally, a proper categorization leads to an appropriate response and an appropriate response favors the individual who made it. This procedure is the result of a prior knowledge that has been obtained from various experiences in life. The perceptual and cognitive strategies that are involved in category learning are one of the most important concepts in cognitive sciences and many studies are focused on understanding the underlying mechanisms of categorization. Most cognitive theorists consider that in humans different category structures require multiple computational processes which are activated by at least two systems of learning with each system related to different brain activity (Maddox et al. 2002; Maddox and Ashby 2004; Hammer et al. 2010). One is the explicit system and is mediated from Rule-Based (RB) learning while the other system is implicit and is responsible of Information Integration (II) learning (Maddox et al. 2004).

Before analyzing the function of each learning system, it is essential to describe first the COVIS model (competition between Verbal and Implicit systems). COVIS is a neurobiological theory based on human studies that describes the competition between the explicit, hypothesis system and the implicit, procedural system which interact closely by neural substrates (Rabi et al. 2015; Helie et al. 2012; Schnyer et al. 2009; Maddox and Ashby 2004). These terms are different from (RB) and (II) as the first describes the methods-strategies of learning that is used by an individual and the second the structure of a categorization task. This distinction is important as it is possible an explicit, hypothesis system to be used in an (II) task (Maddox et al. 2010). This may happen if the participant is taking into account only one single dimension of the properties of the stimuli presented, while the categorization task demands multiple dimensions to be perceived in a pre decisional level (Spiering and Ashby 2008). Multiple dimensions may be involved in a stimuli at once (e.g. frequency/duration) or only one single dimension (e.g. duration). It has been shown, that in visual category experiments, participants used only one single dimension to reach optimal solution (Feldman 2000), whereas trial by trial feedback was most of the time necessary to motivate them to use multiple dimensions (Ashby et al. 1998).

In order to assign different objects in the same group, participants must spot the category-relevant properties of these objects and separate the irrelevant ones (Perry and Lupyan 2014). For example, all blue objects belong to category (A) and all red in category (B). In this case, color is the selective dimension that separates one category from another. Discrimination of these properties and sorting of new groups can be done by using either the explicit, hypothesis system or the implicit, procedural system. Organisms may use both systems equally or it is possible that one system dominates the other. If only one system dominates during category learning, then the cognitive system is considered to be dimensionally polarized (Smith et al. 2012) although some studies suggest that there is at least some degree of interaction between the two systems (Maddox et al. 2010). Categorization can be considered as not dimensionally polarized if both (RB) and (II) can be learned to the same degree and in the same frequency (Smith et al. 2012).

So far, there are studies that have already tested which methods of learning humans, primates and birds are using during categorization tasks with visual or auditory stimuli. When animals are tested in category learning experiments, the identification of the cognitive process is applied by a correlation between the structure of the categorization task and the performance of the individual. In this way, the representation and the comparison between (RB) and (II) performance may indicate a tendency of the individual of using more or less a specific cognitive system.

## The explicit learning system

The explicit, hypothesis-test system dominates on (RB) strcture, and learning occurs by a conscious reasoning process where the rule that maximizes accuracy (i.e., optimal solution) can be easily verbalized (e.g. long/short duration or high/low frequency), (Ashby et al. 2002; Spiering and Ashby 2008; Schnyer et al. 2009; Helie et al. 2012). Working memory and executive attention can be used at the same time to test or replace the hypothesis and lead to possible solutions (Smith et al. 2012). It has been shown that (RB) performance was reduced when the participants were involved in work that requires working memory and executive attention simultaneously with the categorization task (Mida and Rabi 2015). Moreover, the replacement of a rule with a new one is possible depending on feedback signals (Maddox et al. 2004). Indeed, a negative feedback can be used as guidance to change rule or even cognitive strategy (Schnyer et al. 2009). (RB) requires only one single dimension that must be discovered but also multiple dimensions could be relevant, for example (short-red objects as category (A) and long-blue objects as category (B)). In this case (RB) structure demands multiple dimensions that must be taken into account but still the procedure of learning follows a general rule. According to Ashby, in most rule based category trials only one stimulus dimension is relevant, and the observer’s task is to discover this relevant dimension and then to map the different dimensional values to the relevant categories (Ashby et al. 2002; Hélie et al. 2012). The (RB) structure that has been used in this study is displayed in (Fig. 1).

**Fig. 1.**
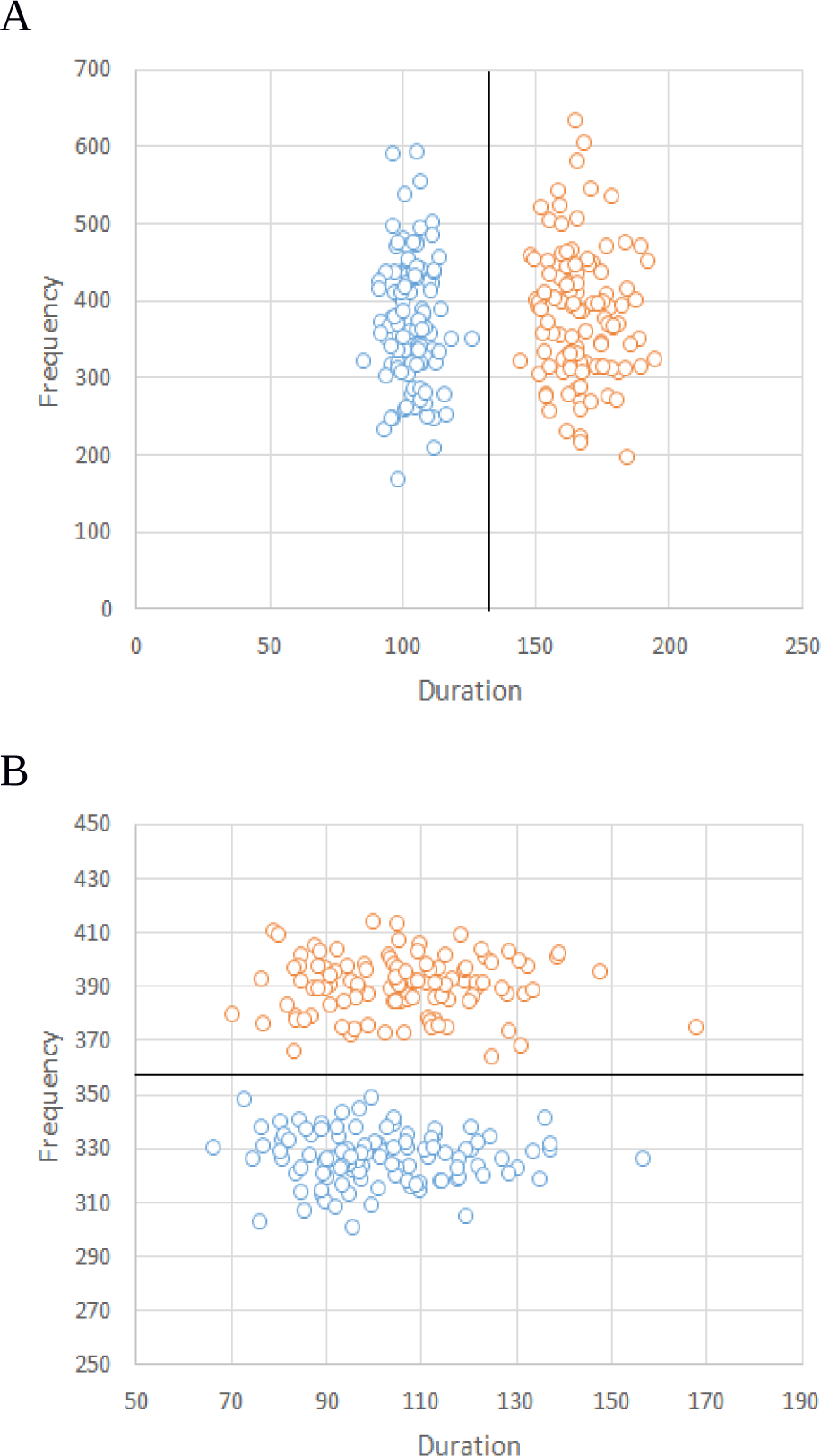
Each circle represents one acoustic unit (Dutch vowel) the blue circles belong to category A while red circles belong to category B. In fig. 1A the relevant dimension is duration. Maximum performance can be achieved if the participant set as a discrimination criterion the black line between the blue circles and the red circles. On the other hand, in fig. 1B the relevant dimension is frequency. In this case participants have to ignore duration and focus on first format frequency. The black line between the two categories as well as fig. 1A displays the way to optimal solution. (Frequency in Hz, Duration in ms).

Different neural mechanisms are activated in brain in (RB) and (II) learning. The prefrontal cortex seems to have an important role in the operation of the explicit, hypothesis-test system (Maddox et al. 2010). The prefrontal cortex is part of the frontal lobe which is located on the front part of the mammalian brain and is responsible of the human ability to plan and follow complex rules (DeYoung et al. 2009) hence it regulates partially, the effective use of feedback (Maddox et al. 2010). Feedback delay or its representation prior to stimuli does not affect rule learning but this system learns faster under full use of feedback (Maddox et al. 2008).

The explicit, hypothesis-test system is not only associated with the prefrontal cortex but also with hippocampus whose activity depends on the requirements of the categorization task. Hippocampus activity has more involvement when grouping stimuli into two different categories than grouping stimuli inside the same category (Seger and Peterson 2013). More parts of the brain are included on rule-based learning; some of them are the medial temporal cortex, the anterior cingulated cortex, and the head of the caudate nucleus (Ashby and Maddox 2010). There is a close interaction between these brain components rather than an independent activity as there are several parameters that determine which part of the brain activates more during (RB) learning.

## The implicit learning system

The implicit, procedural learning system dominates on (II) tasks and its strategy relies on integration of two or more dimensions of stimuli in a pre decisional level (Maddox et al. 2010; Spiering and Ashby 2008; Maddox et al. 2004). Unlike the explicit system, the participants are unable to describe verbally the method they are using and they lack consciousness to the reasons of their behavioral responses (Smith et al. 2012; Helie et al. 2012). Manipulations of feedback such as delay for a few seconds impair the process of implicit learning (Smith et al. 2012). Furthermore, in contrast with the explicit system, working memory and executive attention does not affect the outcome of the performance during categorization (Soto et al. 2013). (II) Structure is formed in a way that maximum performance can be discovered without following a rule. Properties of the stimuli are perceived as a whole and not as separate components, for example, the image of a car is processed as one object with multiple dimensions and not as separate subset dimensions that form a superior entity. (Fig. displays (II) Structure that used in this study.

**Fig. 2.**
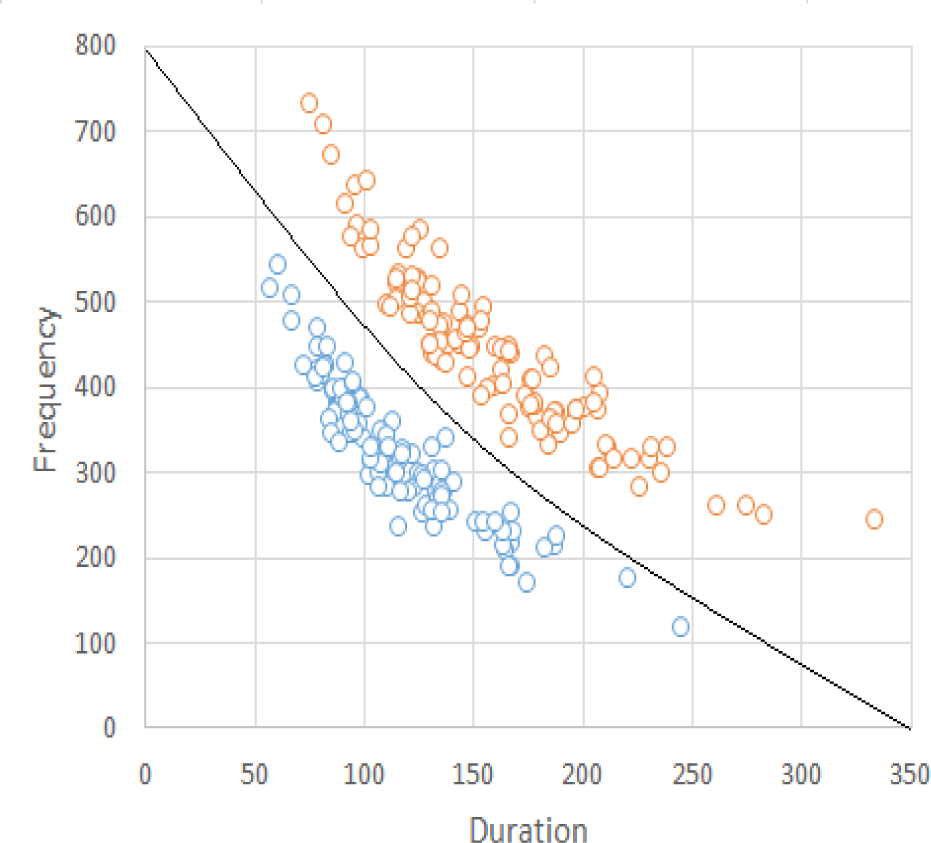
Blue and red circles represent category A and B respectively. In contrast with (RB), participants should perceive the acoustic stimuli (Dutch vowel) as a multidimensional unit and base their response relying in two dimensions (both frequency and duration). The black curve between the two categories determines the discrimination boundary that leads to best performance. A categorization strategy that relies in only one single dimension would lead to 50% correct responses and cannot be considered as a learning effect.

Information-integration learning is highly depended on striatum, a subcortical part of vertebrate forebrain (Maddox et al. 2010). Parkinson’s disease patients (PD) with impaired striatum function show poor procedural learning. However, some studies imply that (PD) patients have low accuracy on both (RB) and (II) tasks indicating that striatum might have a multiple role in category learning (Hammer et al. 2010). Consequently, some components of basal ganglia have a major role in the implicit, procedural system. Basal ganglia is located at the base of forebrain and interacts with posterior perceptual regions during (II) learning (Nomura et al. 2006). Structure of basal involves the putamen, globus pallidus and the caudate nucleus. According to COVIS model, (II) tasks activate visual cortical areas and the posterior caudate nucleus (Grimm and Maddox 2013). Just like explicit system, the procedural system recruits a number of different brain areas that interact as a system and not separately.

## Beyond human category learning

Research on cognition systems using categorization tasks has progressively lead many studies to use the same methods that are used in humans also in primates and birds. Generally, the progress of learning during a categorization task indicates which cognitive system the subject is using more. For example, the abrupt increase in the performance in a (RB) structure implies that the subject might had discovered a rule and discriminated easily between categories. This process is correlated to the explicit, hypothesis-test system. According to Smith, humans, rhesus macaques, and capuchin monkeys had lower performance on (II) than (RB) when tested in visual category tasks. All species struggled to integrate information across two stimulus dimensions, while their accuracy improved significantly on tasks that required only one single dimension, showing that process of learning occurs faster and easier using the explicit system instead of the procedural learning system (Smith et al. 2011). The ability of using multiple strategies during category learning suggests that the individual conceivably have more than one cognitive system. In contrast, pigeons showed no tendency upon a cognitive strategy and solved rule base and information integration tasks equally without significant difference in overall performance (Smith et al. 2011). More studies support the idea that pigeons may have a non analytic way in category learning implying that the utility of multiple cognitive systems may be restricted to primates (Berg et al. 2013; Smith et al. 2010).

This research focuses on the Zebra Finch, a songbird species that belong to the Estrildidae family. Zebra finches are able to identify their conspecifics (individuals or groups) relying on the songs that they are exposed to, making them that way a suitable model for studying the acoustic patterns that may exist during categorization (Nagel et al. 2010). Moreover, they represent many similarities with humans, particularly in vocal and auditory learning. Discrimination of vocal and auditory learning is important since the first describes the way to memorize a sound and then the ability to imitate it or even modify it, whereas the second refers to the process in which learning occurs from sounds heard (Bolhuis and Everaert 2013). Only three avian and three mammalian orders are able of vocal learning, (songbirds, hummingbirds, parrots) and (humans, bats and cetaceans) respectively. Since there are distant relatives in the three avian orders that are capable of imitation and improvisation of sounds, it is suggested that they have evolved vocal learning independently (Jarvis 2006). Both humans and Zebra finches learn through auditory perception which later serves to guide vocal production (Weisman et al. 2014) and even from an early age they both imitate sounds from their surroundings and especially from their parents. In this early period of life learning occurs faster and easier (Bolhuis and Everaert 2013). Auditory feedback is necessary to maintain and develop vocal production as songbirds use their own vocalizations as a template which progressively alters in a “crystallized” song which is an accurate copy from their parent’s song. Parallels that have been found between human and songbird’s brain might indicate that vocal learning in both species relies on similar brain mechanisms. Furthermore, examples of deep homology also exist, FoxP2 gene expression is most studied as it has a major role in vocal production, vocal learning and coordination in both species (Bolhuis & Everaert 2013).

## Present study and testing hypothesis

There is a considerable amount of literature on parallels between human speech and birdsong but less are known about the underlying cognitive mechanisms that enhance the process of auditory learning. In this research we analyze and compare data between humans and Zebra finches that are tested in auditory category tasks with equivalent methods and identical stimuli. In particular, we used a modification of (Goudbeek et al. 2008) stimuli in order to investigate the methods of learning.

In Goudbeek’s experiment Castilian Spanish and American English students participated in speech category learning tasks in which they had to properly discriminate and categorize synthesized Dutch vowels /Y/, /y/, and /ø/. Spanish listeners learned to distinguish between /Y/ and /ø/ or /y/ and /Y/, with or without feedback. In both conditions (/Y/-/ø/ and /y/-/Y/), vowels could be discriminated based on a single dimension (duration and first formant frequency, respectively). American listeners also had to categorize between /Y/ and /ø/ with supervision or they had to classify /y/ and /ø/ which differ both in duration and in first format frequency. Spanish listeners learned to differentiate between the Dutch vowels easier, when trials based on first format frequency than on duration and their performance reduced significantly when they tested without trial-by-trial feedback in both cases. In contrast, American listeners were better able than the Spanish listeners to acquire the duration-based contrast. However their performance decreased when they tested on multidimensional tasks with trial-by-trial feedback (Goudbeek et al. 2008).

We repeat Goudbeek’s experiment in Zebra finches and in Greek participants who had no prior knowledge of Dutch. We examine which learning strategies these species will operate and compare data to unveil differences between avian and primate cognition. Particularly, in this study, Dutch vowels differ depending on three conditions. In condition (1) vowels differ in duration only (/ø/-/Y/), in condition (2) in first formant frequency only (/ Y/-/y/), and both first formant frequency and duration (/ø/-/y/) in condition (3). In condition (1) and (2) proper categorization of the stimuli is feasible if the response relies in one single dimension (duration or first format frequency respectively for each condition), while both dimensions must analyzed simultaneously in condition (3). Considering that (/ø/-/y/) differ in multiple dimensions, condition (3) is related to (II) strategy while condition (1) (/ø/-/Y/) and condition (2) (/Y/-/y/) corresponds to (RB) strategy as they involve only one single dimension. For details of the acoustic stimuli used in this study see (Table 1).

**Table 1.**
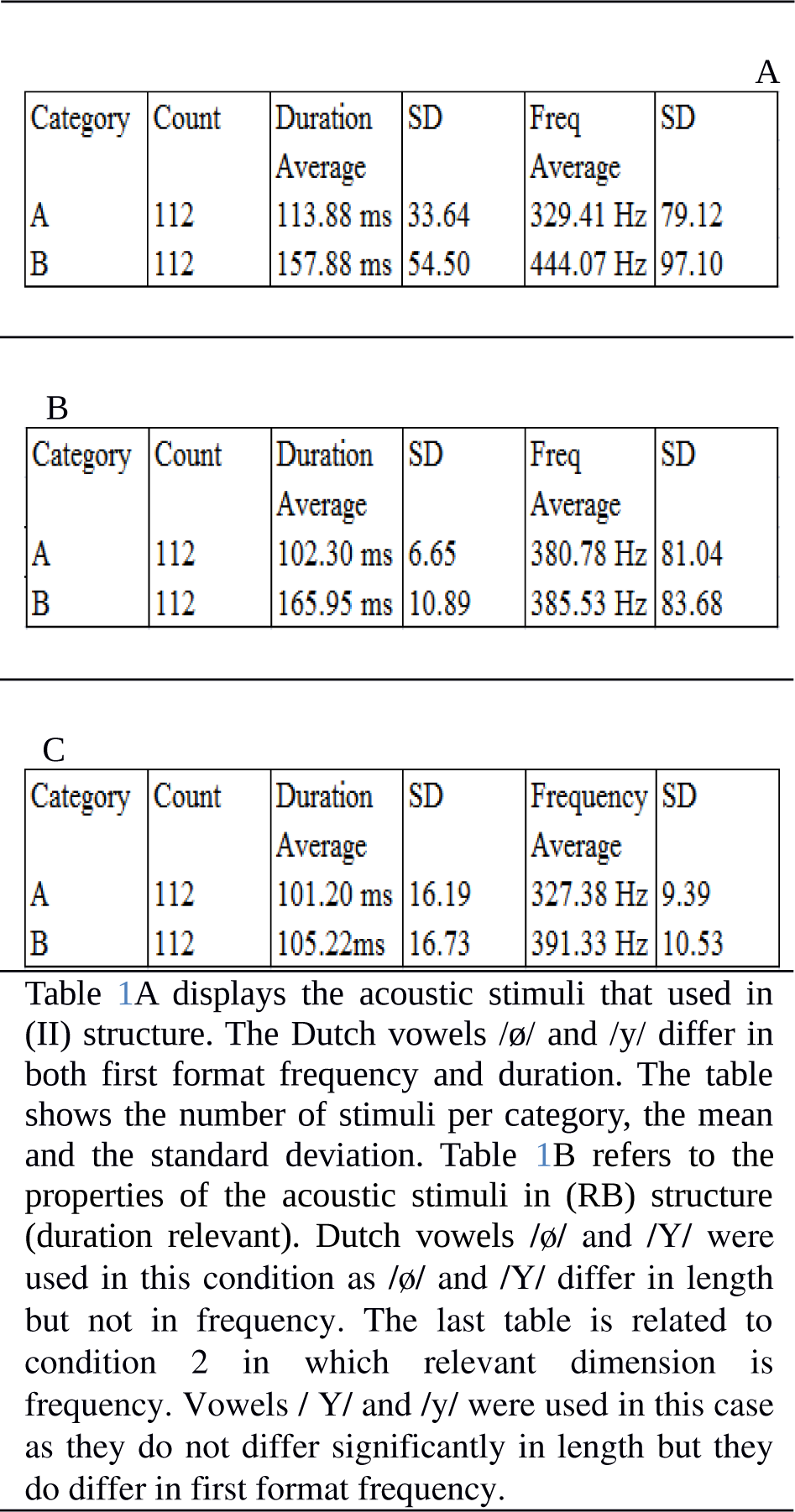
Properties of Dutch vowels “/Y/”, “/y/”, and /ø/ used in the experiment

Two Zebra fiches and four Greeks were tested in each condition. Feedback was given for correct and incorrect responses. When Zebra finches were tested, we divided the experiment in two phases that proceed in sequence: *shaping phase* and *learning phase*, instead with humans that were tested only in *learning phase*. The *learning phase* consisted of 224 stimuli (2 categories x 112 stimuli per category). Zebra finches repeated the test until they reached (75%) of correct responses for three consecutive days while humans were tested in only two repetitions, (448 stimuli overall). Learning was analyzed based on % correct responses and d-prime (Goudbeek et al. 2008).

Since pigeons learned equally both (RB) and (II), our hypothesis is that songbirds do the same providing that way more data on the assumption that the utility of multiple learning strategies are indeed a privilege restricted to primates. On the other hand, due to similarities in auditory and vocal learning that Zebra finches and humans share, simultaneously to our first hypothesis, we do not exclude the chance that both species might use the same methods of learning.

## 2. Methods

### Experiment 1

Twelve Greek adults from Thessaloniki (four in each condition) with no prior knowledge in Dutch language were tested. Data about gender, age, health issues and problems of hearing (if exist) were taken to evaluate if the subject is able of participating to the research. Participants had to react and respond to a specific auditory stimulus and classify it into groups, providing that the subject is capable of discriminating one stimulus from another. The response positions were reversed from subject to subject to avoid any underlying bias toward right or left choice.

Prior to Greek participants we also tested six Dutch adults so as to confirm that the stimuli can be compared to the realistic sound of a Dutch vowel. Greeks on the other hand filled a questioner after the experiment in which they were asked in what degree they perceived the stimuli they heard as vowels. Both Greeks and Dutch reported high proportion of resemblance.

All participants were placed in front of a computer with two boxes on the right and on the left of the screen as response keys and in the middle an image that simulated the representation of a sound. Soundproof headphones (Philips SHP 2000) were provided in order to minimize the effect of external noise. Each time they selected the middle image a Dutch vowel was presented and they had 5 seconds to categorize the vowel either to the right or to the left box. If during that time the participant didn’t respond, the next trial was displayed and the answer was evaluated as “no response”. Feedback was given for correct and incorrect answers. The categorization task was consisted of 448 stimuli (2 categories x 2 repetitions x 112 stimuli per category) (Goudbeek et al. 2008). They were not given any further information except that they had to categorize Dutch vowels and that they would get feedback for each response.

### Results and discussion

First, we divided the trials in experiment in half in order to detect the learning effect that occurred during categorization. Specifically, we set as learning phase 1 the first 224 stimuli and one repetition of these as learning phase 2. A d’ prime analysis was conducted and the percentage correct of each phase were calculated. (Fig. 3) displays the results of each condition in d’ prime and percentage correct.

**Fig. 3.**
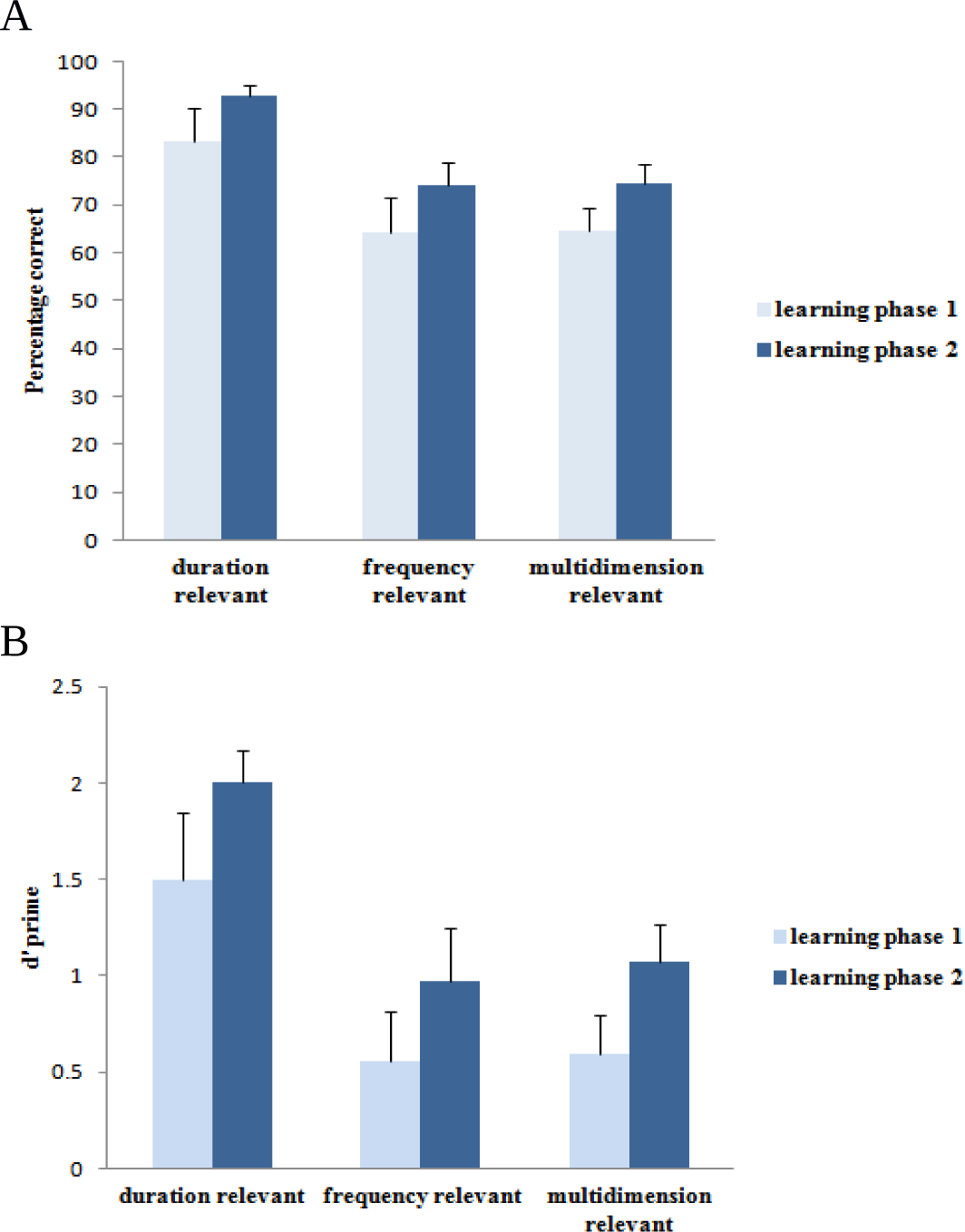
Performance in percentage correct fig. 3A and d’ prime fig. 3B for learning phase 1 and 2 in each condition.

Difference in correct responses rate between learning phase 1 and 2 point out a learning effect but further analysis is necessary to conclude that participants are able to discriminate between categories. For this purpose, we used signal detection model to indentify if participants were able to discriminate between the acoustic stimuli. Accurate discrimination among two stimuli that belong to unrelated categories is evaluated as a “hit” (H) while failure to spot the difference as “false alarm” (F). The variation between the z-transforms of hit and false alarm rate z (H) - z (F), (H) = “hit rate”/”total # responses” (F) = “false alarm rate”/”total # responses” is correlated to the value of d’ prime.

Generally, the results illustrate a better performance in phase 2 than in phase 1. The percentage correct was significantly above chance in phase 2 in all conditions (condition 1 t(3) = 18.19, p<.005, condition 2 t(3) = 5.03, p<.005, condition 3 t(3) = p<.005). D prime ratio was also significantly above zero in all conditions except phase 1 in frequency and multidimensional condition (condition 1 t (3) = 4.86, p<0.05 and t(3) = 11.2, p<.005 in phase 1 and 2 respectively, condition 2 t3 = p<.005, condition 3 t(3) = 5.32, p<.005). Furthermore, an ANOVA analysis was conducted to uncover any significant differences between conditions in phase 1 and 2. The analysis showed that phase 1 had no significant difference in correct percentage between conditions while d’ prime was close to significance (ANOVA: F2, 9 = 4.03, P = 0.056). In contrast with phase 1, our analysis showed significant difference in phase 2 in percentage correct and d’ prime (ANOVA: F 2, 9 = 8, P = 0.01 and F2, 9 = 5.7, P = 0.02). Data revealed also that phase 1 and 2 differed significantly among conditions in correct percentage and d’ prime respectively (ANOVA: F3, 5 = 3.34, P = 0.04 and F3, 5 = 0.81, P = 0.05). Condition 2 did not differ significantly from condition 3. However, both differed with condition 1 where duration was the relevant dimension (conditions 1 and 2, t(4) = 3.43, p<.005, t(4)= 2.86, p<.005 for correct percentage and d’ prime respectively and conditions 1 and 3, t(5) = 4.01, p<.005, t(6) = 3.46, p<.005).

### Experiment 2

Four male Zebra finches were chosen in experiment 2. All of the birds were born and raised in the laboratory of Animal Behavior department in Leiden University. None of them had been used before in previous experiments. They had constant access to water and grit and no problems of health reported during the experiment. The procedure of the experiment consisted in two phases that proceed in sequence *shaping phase* and *learning phase.* First, each bird was placed individually in a cage in a sound-attenuated room. Above the cage, speakers was installed and presented the auditory stimuli, (songbirds or Dutch vowels) depending on the phase. Three sensors with a LED were set in a line inside the cage. Sensor 2 (s2) and sensor 3 (s3) determined the responses while sensor 1 (s1) which was set in the middle produced the sound stimulus. A modified Skinner box was registering and controlling the responses and the presentation of the stimuli. The Skinner box was connected to LEDs, sensors, speaker, food hatch door and light of the room. Fig. 3 displays the dimensions and the structure of each cage.

**Fig. 3.**
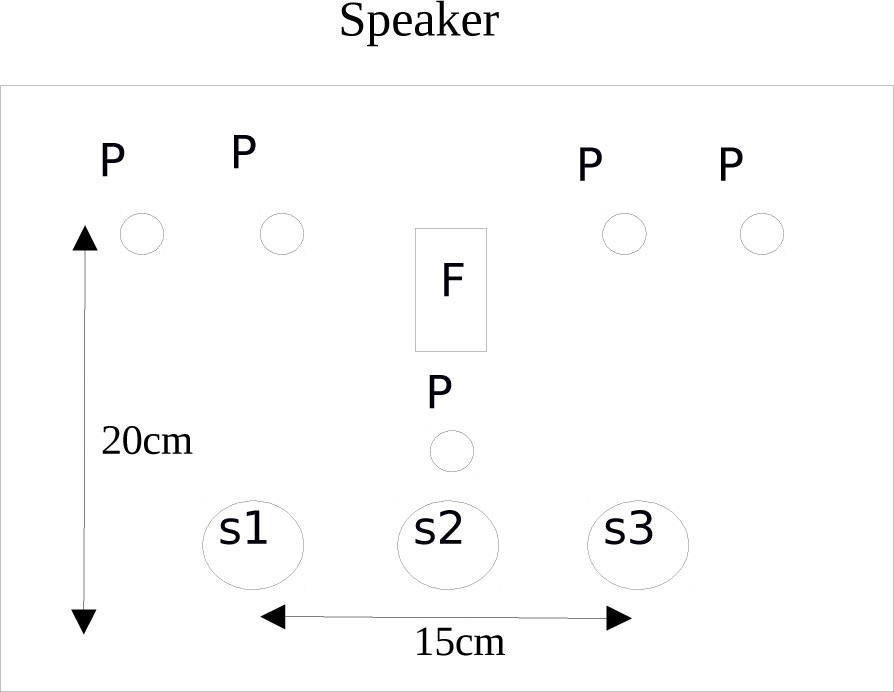
Structure of the experimental cage. The speaker was set above the cage (40cm) and represented sound stimuli when the bird pecked (s1). (s2) and (s3) were placed at equal distance from (s1) (7cm) each. Above (s1) a vertical perch (p) was set and the bird could have access to food hatch door (F).The light was placed above the cage and was regulating day time and night time (07:30 lights on and 20:00 lights off).

### Shaping phase

All Zebra finches were placed individually in an experimental cage for two days to adjust to their new environment. Over these two days, the food hatch door remained open. After this period, the shaping phase begun and was divided in four stages. In stage 1, the food hatch door was closed and LEDs of all sensors was on. Each time a bird pecked (s1), birdsong 1 (b1) was presented and the food hatch door opened for 15 seconds. Sensor 2 and (s3), produced also (b1) but the food hatch door remained closed. This stage continued until the bird reached approximately 200 pecks of (s1). In stage 2, only (s1) was on and produced (b1) after a peck. In order to have access to food, the bird had to choose between (s2) and (s3) within 15 seconds. Sensor (2) was set as a reward (15 seconds food hatch door open) and (s3) as a punishment (1-5 second lights off). We waited until birds reached above 75% percent of correct responses for at least 24 hours before we proceed in the next stage. Stage 3, differed only in the acoustic stimuli (birdsong 2 (b2) instead of (b1)) and feedback from (s2) and (s3) was reversed. Same as stage 2 the process continued until 75% of correct responses. In the last stage of shaping phase, (s1) was playing either (b1) or (b2) (ratio 50%). (s2) was set as a correct response to (b1) whereas (s3) as punishment and vice versa when sound stimuli was (b2). Punishment increased to 15 seconds and reward decreased to 10 seconds. In contrast with previous stages, the percentage correct had to be more than 75 % for three consecutive days before we proceed to learning phase. The completion of stage 4 was set as the decisive factor that confirmed that the birds categorizing according to the sounds they heard and not randomly.

### Learning phase

During this phase, speaker presented only Dutch vowels that differ in duration, frequency or both depending on the condition. Four birds were tested on 448 stimuli (2 categories x 112 stimuli per category). Number of repetitions depended on bird’s skill to reach 75% correct responses for at least three consecutive days (learning criterion). After a peck on (s1) a Dutch vowel was represented and then the bird had to categorize the stimuli either to (s2) or (s3). Feedback was given for correct and incorrect responses (10 seconds food hatch door open - 15 seconds lights off). Time limit of each trial was 15 seconds. If there was no response after 15 seconds the stimuli was repeated until bird responded.

### Results and discussion

Two Zebra finches were tested in frequency condition, one in duration and one in multidimensional condition. A bird passed away during the experiment due to software malfunction and its data did not included in the results. Fig. 4 shows the results of each bird in each condition.

**Fig. 4.**
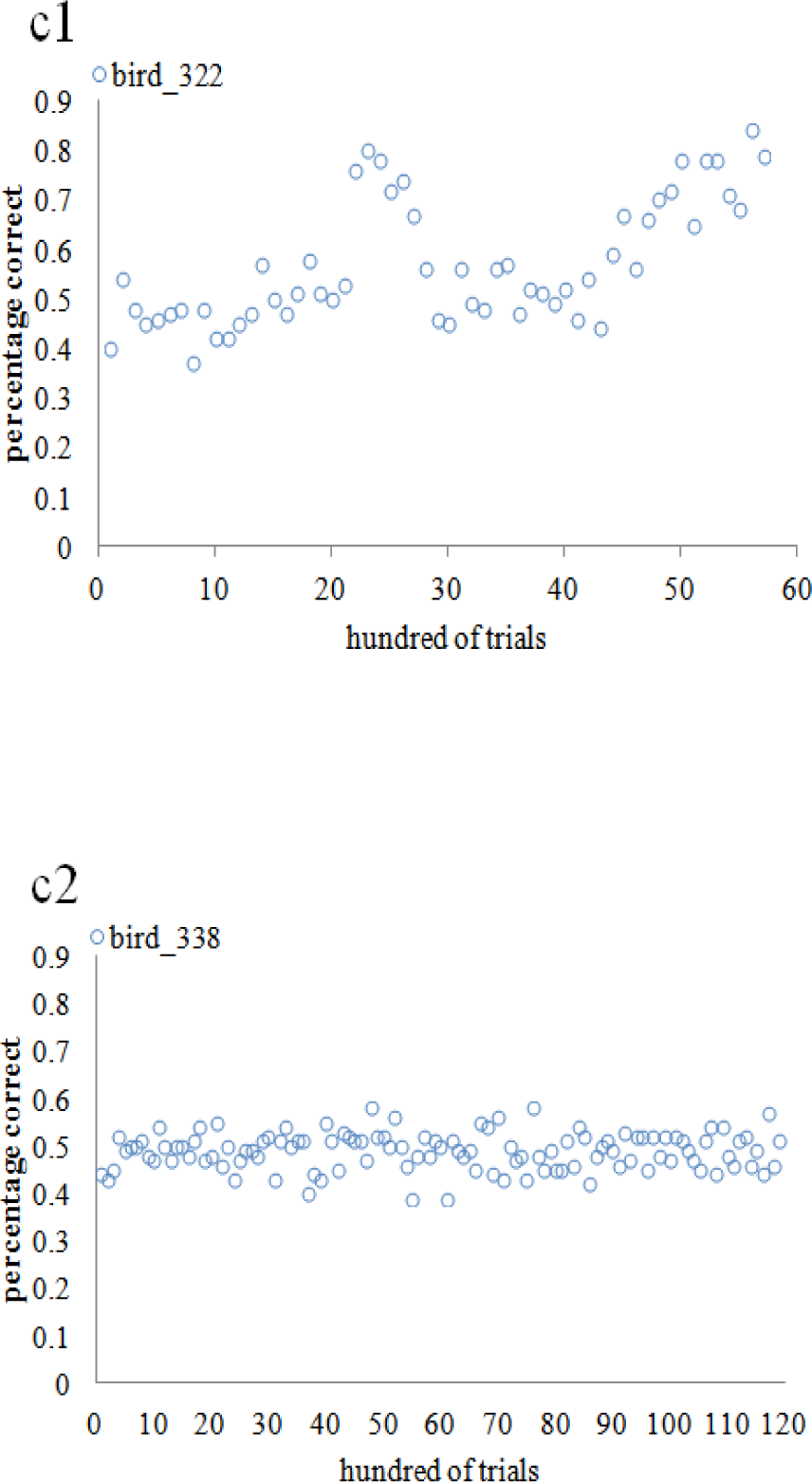
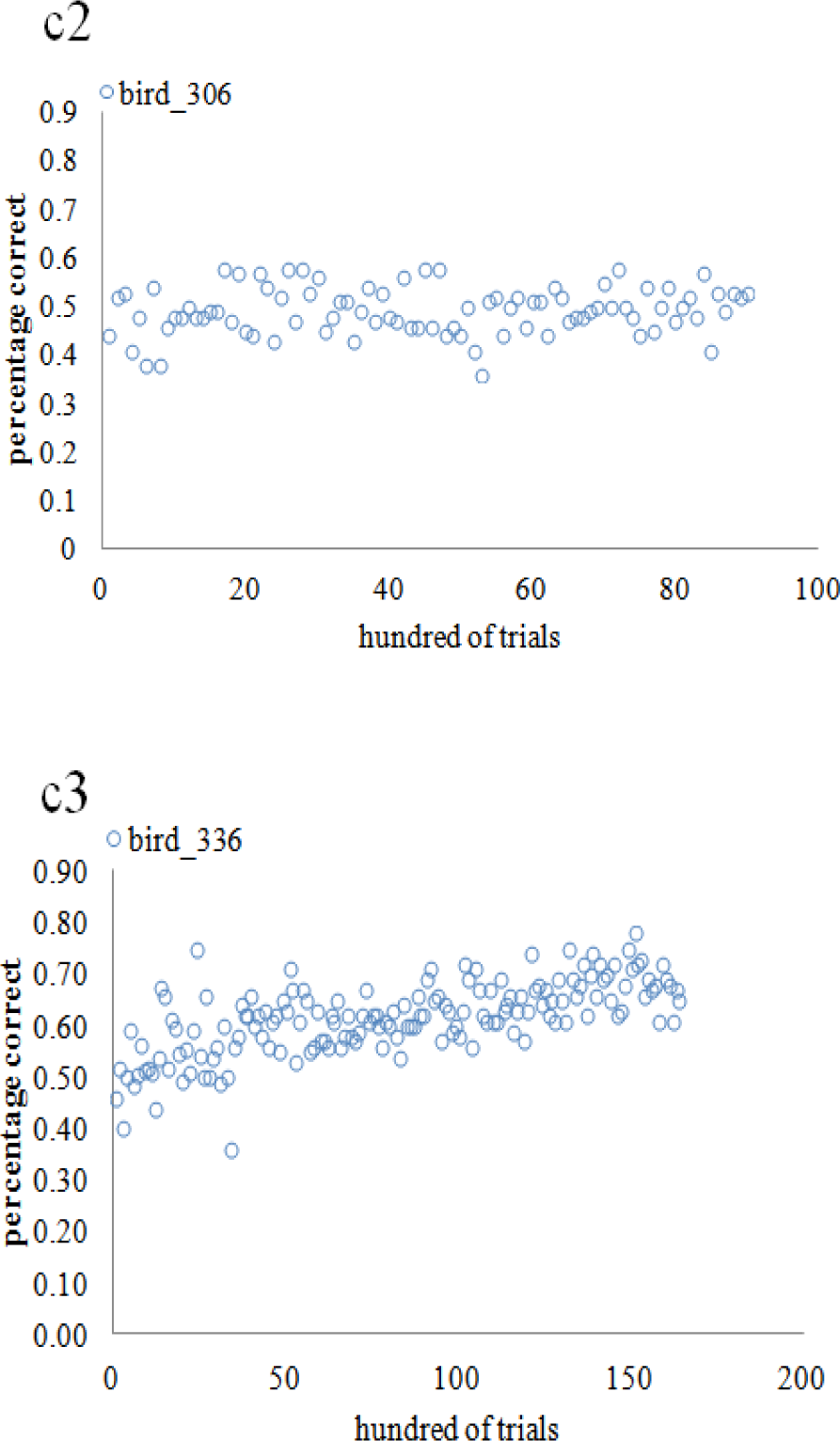
Percentage correct for each bird in c1, c2 and c3. Each blue circle represent a hundred of trials, the first scatter plot shows the learning effect of bird_322 in duration relevant condition while the second and third scatter plot the performance of bird_338 and bird_306 in frequency relevant condition. The last scatter plot represents the accuracy of bird_336 that tested in the multidimensional condition. In the multidimensional condition an error occurred during the experiment and for 2000 trials approximately the same acoustic stimuli was repeated until bird_336 gave a correct response.

In contrast with pigeons that learned (RB) and (II) structures equally (Smith et al. Zebra finches showed a discrepancy in the way of learning different category structures. The acquisition of (RB) structure in duration relevant condition occurred abruptly and performance reached 75% of correct responses in less than 6000 trials.

Overall, bird_322 required 23 days to reach our criterion (13 April to 6 May). Contrariwise, the acquisition of the related category occurred progressively and far more slowly in the multidimensional condition and bird_336 needed 150000 trials approximately until its performance reached 70% of correct responses. In total, bird_336 needed more than a month to complete the experiment. In condition 2 bird_338 and bird_306 performed at chance during all time of the experiment. We investigated further the procedure of learning in condition 2 and we examined in detail the responses of each bird. Fig. 5 shows the response tendency of (s2) and (s3).

**Fig. 5.**
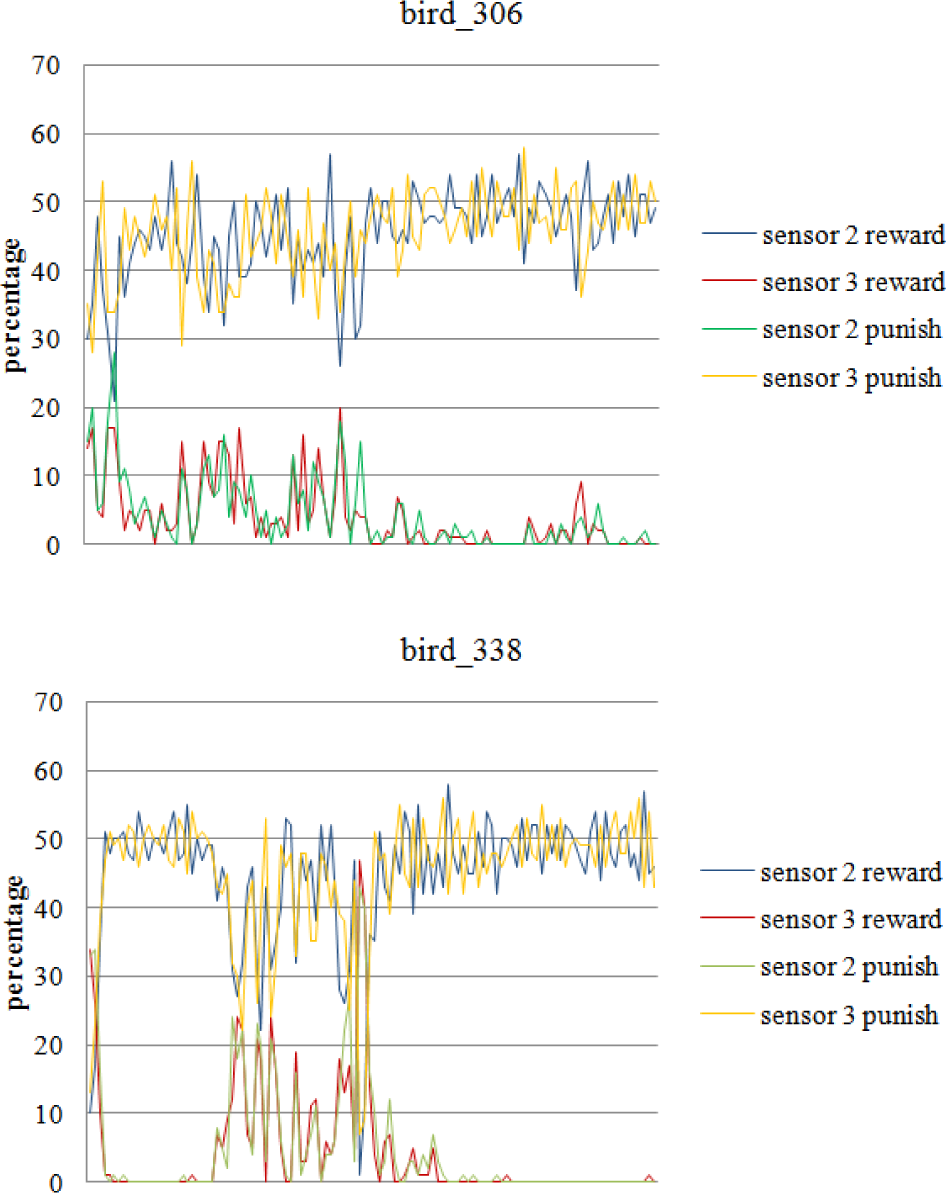
The percentage (number of pecks) for each sensor during the experiment. Colored lines represent the ratio of reward and punishment for (s2) and (s3) respectively.

It is clear, that both birds showed a strong tendency on pecking sensor 2 and ignored the punishment of sensor 3 during the experiment. The reasons of this behavior are still unidentified but this result indicates that none of the birds made a realistic effort of learning to discriminate between the two categories. Therefore, we are not able to conclude that Zebra finches actually failed to categorize /Y/ and /y/.

Furthermore, we examined the learning strategy of bird_336 in the multidimensional condition and an analysis in level of individual stimuli was made. Particularly, we calculated the total number of trials and we divided the experiment in 4 equal blocks with X number of trials per block. Initially, the total number of correct and incorrect responses was measured for every sound and every block. For each block the response to the sound stimuli was registered either as “correct” or “incorrect”. For example, if the bird had more correct responses than incorrect on sound_001 this sound was registered as “correct” for the relative block. Fig. 6 shows the learning process in the information integration structure.

**Fig. 6.**
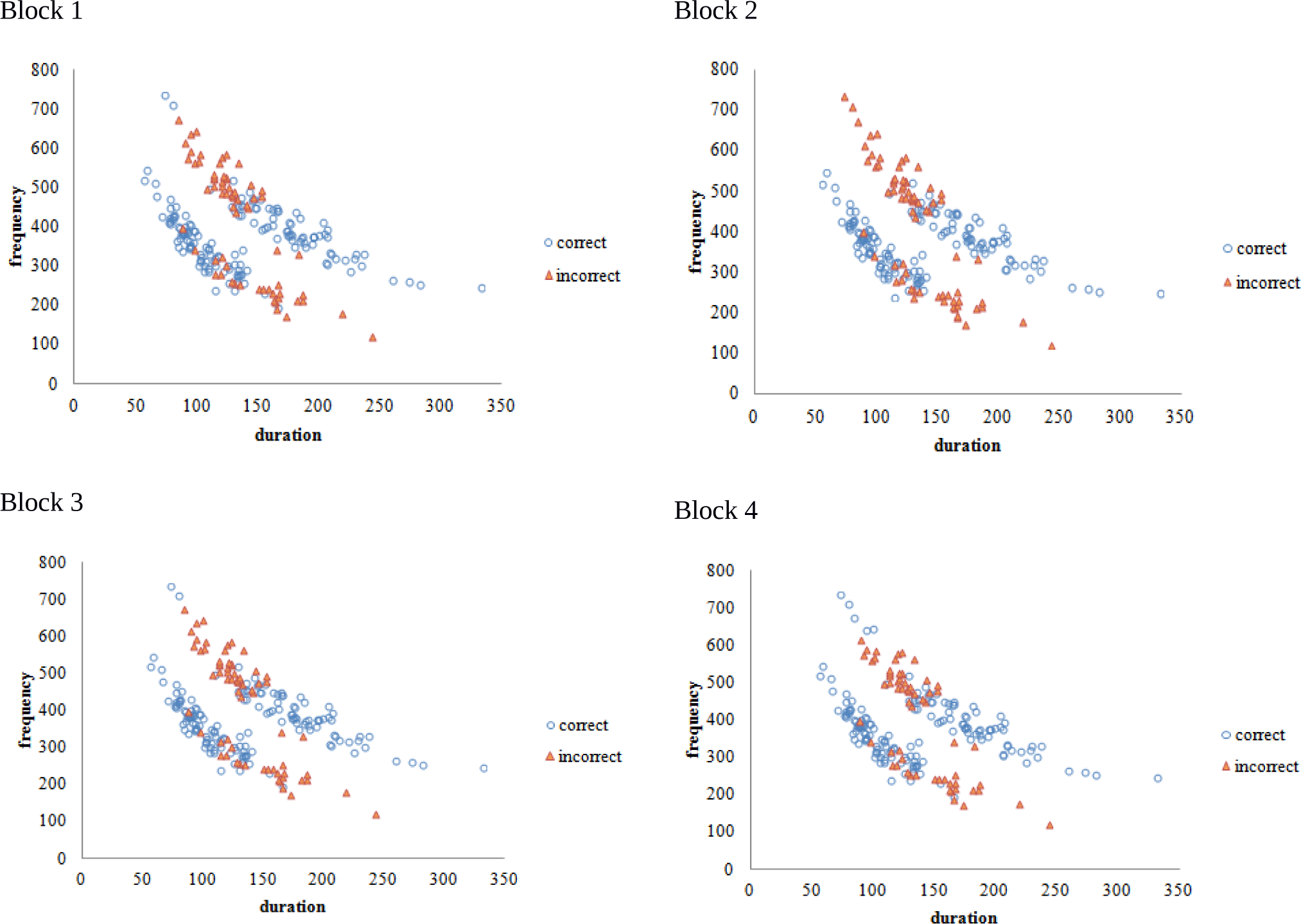
Each block is related to 3.371 trials. Blue circles represent the individual exemplars that bird_336 categorized properly while the triangles are associated to the exemplars that categorization failed.

Bird_336 applied the same strategy during the whole experiment and only minor changes occurred in the procedure of learning. Most of the stimuli with frequency values below 230 Hz were mistakenly characterized as category B. At the same time, bird failed to categorize half of the stimuli with duration values from 100 ms to 150 ms and frequency values from 430 Hz to 600 Hz. The descriptive analysis shows that bird_336 didn’t use any decision boundary that would be used in a rule based structure. The graph indicates that bird_336 was able to perceive the basic structure of the categorization task although it didn’t succeed to reach our criterion.

Finally, to examine further the learning process in both species we analyzed the learning curves during experiment 1 and 2 in duration relevant and in multidimensional condition. Frequency condition was not included because Zebra Finches didn’t accomplish to learn this condition. Both species expressed high sensitivity on duration during categorization and learnt faster and to higher percentage condition 1 than condition 3. Fig. 7 shows the progress of learning of four Greek adults and two Zebra finches in condition 1 and 3.

**Fig. 7.**
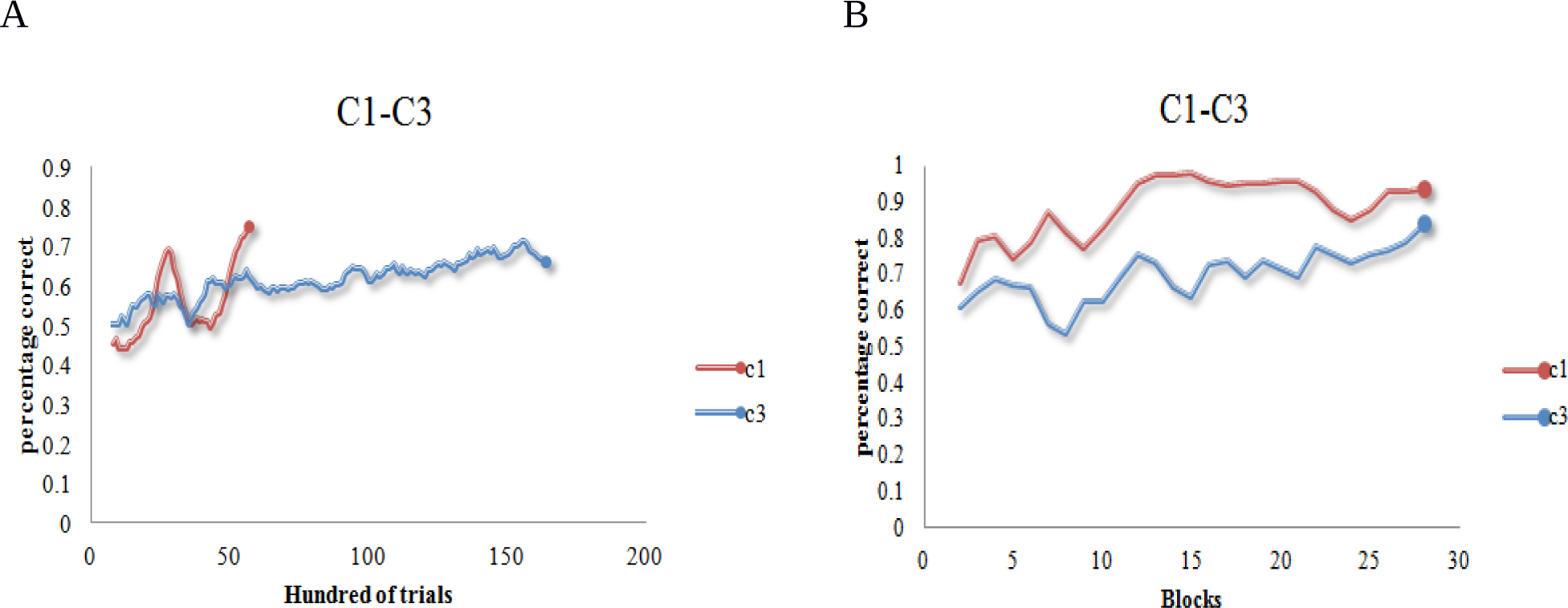
Graph A represents the progress of learning of bird_336 and bird_322. Red line is related to the duration relevant condition (c1) and blue to the multidimensional condition (c3). Same in graph B it is displayed the learning effect for the Greek listeners. Each block in graph B is consisted of 16 trials. In both cases performance was higher in the one-dimensional categorization task and the discrimination criterion reached in less time than in multidimensional condition.

### General discussion

The aim of this research was to investigate the cognitive differences between human and avian cognition by testing Greek adults and Zebra finches in speech categorization tasks, using the same stimuli and equivalent methods. Twelve Greeks and four Zebra finches were tested in three different conditions. Proper categorization in each condition was possible if the subject could discover the relevant properties of the acoustic stimuli and distinguish one category from another. In experiment 1 both d’ prime and correct percentage were above zero and chance respectively in phase 2 of all conditions. Participants were able to categorize the Dutch vowels and all subjects exceeded 75% correct responses in condition 1. Greek participants revealed high sensitivity on duration during the categorization task and gave less attention to frequency in contrast with the Spanish listeners on (Goudbeek’s et al. 2008) experiment. However, our results in multidimensional condition are comparable with Goudbeek who also tested American listeners with supervision in Dutch vowels that differed in two dimensions simultaneously. The faster acquisition of the related categories and the higher performance that Greek listeners achieved in condition 1 than in the other two conditions imply that they trust their responses more in duration than in frequency. The rapid increase that occurred during condition 1 from 75% to 100% which remained until the end of the experiment indicates that the participants might have discovered a rule at some point during the experiment. The fact that the participants presumably detected a rule in a (RB) structure corresponds with the idea that they used a learning method that is associated with the explicit hypothesis-test system. On the contrary, the gradual increase in performance in condition 3 that was organized by an (II) structure, illustrate a diffuse way of learning that can be related to the implicit procedural system. Our results are comparable with (Smith et al. 2012) and support the hypothesis that humans have multiple systems of categorization plus that each system develops in a different way. In condition 2 that frequency was the relevant dimension, Greek listeners had the same proportion of correct responses as in condition 3 in both phases. The signal detection analysis showed that the discrimination ability in these two conditions was similar.

Experiment 2 was conducted in two phases (shaping and learning). Bird_338 and bird_306 were tested in frequency relevant condition but they did not manage to distinguish between categories. Further analysis showed that no effort of learning observed during the experiment as both birds were constantly pecked only one out of two sensors and ignored the punishment that they got for the incorrect responses. More research needs to be done in order to understand the reasons for this behavior and to evaluate if birds were able to discriminate between /Y/ and /y/. However, in the other conditions Zebra finches managed to spot the differences between the acoustic stimuli and learnt to categorize properly the Dutch vowels. Interestingly, the process of learning differed notably in condition 1 and 3. Bird_322 learnt condition 1 faster and to higher terminal performance than bird_336 in condition 3. The abrupt increase from 50% to 80% correct responses indicate that bird_322 might have discovered a rule and solved the categorization task faster than bird_336 whose performance increased gradually. No more than 7.000 trials needed for bird_322 to reach our criterion while bird_336 performance reached close to 75% correct responses in 160.000 trials approximately. Further analysis in level of individual exemplars revealed that bird_336 followed a specific strategy that corresponds more to an implicit procedural learning system than to an explicit hypothesis-test system. The fact that bird_336 probably solved the (II) structure using the implicit procedural learning system speculates that Zebra finches are actually capable of integrating both dimensions at the same time in order to categorize properly the acoustic stimuli plus that they might have more than one category learning system.

Our results showed that one Zebra Finch were able to acquire the (RB) task faster and easier than (II). Assuming that this hypothesis is accurate, then the utility of multiple systems of categorization might not be restricted to primates as (Smith et al. 2011) suggest. Due to our limited sample size, it is hard to state that they have more than one cognitive system and that they learn speech categories just like humans do. For this reason, further research needs to be done to unveil the differences between avian and human cognition. Our results although suggest that Zebra finches probably utilize different methods of learning depending on the structure of the categorization task that they are tested. An important question that emerges out of this result is why pigeons do not use multiple systems of learning but Zebra finches probably do. Zebra finches are able of imitation and improvisation of sounds just like humans. Both species have also a complex system of communication that is based in specific structure and rules. On the other hand, pigeons do not have this privilege. The acquisition of more than one cognitive system is possibly an essential factor that has contributed to the development of this ability.

## References

Ashby, F.G., Alfonso-Reese, L. a, Turken, a U., Waldron, E.M., 1998. A neuropsychological theory of multiple systems in category learning. Psychol. Rev. 105, 442–481. doi:10.1037/0033-295X.105.3.442

Ashby, F.G., Maddox, W.T., 2011. Human category learning 2.0. Ann. N. Y. Acad. Sci. 1224, 147–161. doi:10.1111/j.1749-6632.2010.05874.x

Bolhuis, J.J,. Everaert, M., Berwick,. R.C, Chomsky,. N. 2013. Birdsong Speech and Language: Exploring the Evolution of Mind and Brain.

Berg, M.E., Ward, M.D., Dai, Z., Arantes, J., Grace, R.C., 2014. Comparing performance of humans and pigeons in rule-based visual categorization tasks. Learn. Motiv. 45, 44–58. doi:10.1016/j.lmot.2013.11.001

DeYoung, C.G., Hirsh, J.B., Shane, M.S., Papademetris, X., Rajeevan, N., Gray, J.R., 2010. Testing predictions from personality neuroscience. Brain structure and the big five. Psychol. Sci. a J. Am. Psychol. Soc. / APS 21, 820–828. doi:10.1177/0956797610370159

Feldman, J., 2000. Minimization of Boolean complexity in human concept learning. Nature 407, 630–633. doi:10.1038/35036586

Goudbeek, M., Cutler, A., Smits, R., 2008. Supervised and unsupervised learning of multidimensionally varying non-native speech categories. Speech Commun. 50, 109–125. doi:10.1016/j.specom.2007.07.003

Grimm, L.R., Maddox, W.T., 2013. Differential impact of relevant and irrelevant dimension primes on rule-based and information-integration category learning. Acta Psychol. (Amst). 144, 530–537. doi:10.1016/j.actpsy.2013.09.005

Hammer, R., Brechmann, A., Ohl, F., Weinshall, D., Hochstein, S., 2010. Differential category learning processes: The neural basis of comparison-based learning and induction. Neuroimage 52, 699–709. doi:10.1016/j.neuroimage.2010.03.080

Hélie, S., Paul, E.J., Ashby, F.G., 2012. Simulating the effects of dopamine imbalance on cognition: From positive affect to Parkinson’s disease. Neural Networks 32, 74–85. doi:10.1016/j.neunet.2012.02.033

Jarvis, E.D., 2006. Evolution of brain structures for vocal learning in birds: a synopsis. Ars Zool. Sin. 52, 85–89.

Maddox, W.T., Ashby, F.G., Ing, a D., Pickering, A.D., 2004. Disrupting feedback processing interferes with rule-based but not information-integration category learning. Mem. Cognit. 32, 582–591. doi:10.3758/BF03195849

Maddox, W.T., Love, B.C., Glass, B.D., Filoteo, J.V., 2008. When more is less: Feedback effects in perceptual category learning. Cognition 108, 578–589. doi:10.1016/j.cognition.2008.03.010

Maddox, W.T., Molis, M.R., Diehl, R.L., 2002. Generalizing a neuropsychological model of visual categorization to auditory categorization of vowels. Percept. Psychophys. 64, 584–597. doi:10.3758/BF03194728

Maddox, W.T., Pacheco, J., Reeves, M., Zhu, B., Schnyer, D.M., 2010. Rule-based and information-integration category learning in normal aging. Neuropsychologia 48, 2998–3008. doi:10.1016/j.neuropsychologia.2010.06.008

Minda, J.P., Rabi, R., 2015. Ego depletion interferes with rule-defined category learning but not non-rule-defined category learning. Front. Psychol. 6, 1–9. doi:10.3389/fpsyg.2015.00035

Nagel, K., Kim, G., McLendon, H., Doupe, A., 2011. A bird brain’s view of auditory processing and perception. Hear. Res. 273, 123–133. doi:10.1016/j.heares.2010.08.008

Nomura, E.M., Maddox, W.T., Filoteo, J. V., Ing, a. D., Gitelman, D.R., Parrish, T.B., Mesulam, M.M., Reber, P.J., 2007. Neural correlates of rule-based and information-integration visual category learning. Cereb. Cortex 17, 37–43. doi:10.1093/cercor/bhj122

Perry, L.K., Lupyan, G., 2014. The role of language in multi-dimensional categorization: Evidence from transcranial direct current stimulation and exposure to verbal labels. Brain Lang. 135, 6672. doi:10.1016/j.bandl.2014.05.005

Schnyer, D.M., Maddox, W.T., Ell, S., Davis, S., Pacheco, J., Verfaellie, M., 2009. Prefrontal contributions to rule-based and information-integration category learning. Neuropsychologia 47, 2995–3006. doi:10.1016/j.neuropsychologia.2009.07.011

Seger, C. a., Peterson, E.J., 2013. Categorization=decision making+generalization. Neurosci. Biobehav. Rev. 37, 1487–1200. doi:10.1016/j.neubiorev.2013.03.015

Smith, J.D., Ashby, F.G., Berg, M.E., Murphy, M.S., Spiering, B., Cook, R.G., Grace, R.C., 2011. Pigeons’ categorization may be exclusively nonanalytic. Psychon. Bull. Rev. 18, 414–421. doi:10.3758/s13423-010-0047-8

Smith, J.D., Berg, M.E., Cook, R.G., Murphy, M.S., Crossley, M.J., Boomer, J., Spiering, B., Beran, M.J., Church, B. a., Ashby, F.G., Grace, R.C., 2012. Implicit and explicit categorization: A tale of four species. Neurosci. Biobehav. Rev. 36, 2355–2369. doi:10.1016/j.neubiorev.2012.09.003

Soto, F. a., Waldschmidt, J.G., Helie, S., Ashby, F.G., 2013. Brain activity across the development of automatic categorization: A comparison of categorization tasks using multi-voxel pattern analysis. Neuroimage 71, 284–297. doi:10.1016/j.neuroimage.2013.01.008

Spiering, B.J., Ashby, F.G., 2008. Response processes in information-integration category learning. Neurobiol. Learn. Mem. 90, 330–338. doi:10.1016/j.nlm.2008.04.015

Tschida, K., Mooney, R., 2012. The role of auditory feedback in vocal learning and maintenance. Curr. Opin. Neurobiol. 22, 320–327. doi:10.1016/j.conb.2011.11.006

Weisman, R., Hoeschele, M., Sturdy, C.B., 2014. A comparative analysis of auditory perception in humans and songbirds: A modular approach. Behav. Processes 104, 35–43. doi:10.1016/j.beproc.2014.02.006

